# Transcriptional regulation of the type II fatty acid synthase complex-encoding gene cluster in *Rhodococcus opacus*

**DOI:** 10.1101/2025.09.25.678588

**Authors:** Paulien Geertje Corlijn Leemans, Axelle Van Eupen, Indra Bervoets, Eveline Peeters, Iris Cornet

**Affiliations:** Research Group BioWAVE Industrial Biotechnology, Department of Biochemical and Chemical Engineering, Faculty of Applied Engineering, University of Antwerp, Antwerp, Belgium; Research Group of Microbiology, Department of Bioengineering Sciences, Vrije Universiteit Brussel, Brussels, Belgium

**Keywords:** *Rhodococcus opacus*, transcription factors, regulation, transcriptional architecture, mycolic acids

## Abstract

*Rhodococcus opacus* is an oleaginous actinobacterium with considerable potential for lipid-based bioproduction, as well as for utilising a variety of carbon sources as substrates, including renewable, cost-effective resources. Although its capacity for triacylglycerol accumulation is well established, the regulatory logic that governs its fatty acid and mycolic acid biosynthesis is still poorly understood. Here, we investigated the transcriptional control of the type II fatty acid synthase (FASII) pathway in *R. opacus* PD630, revealing a regulatory architecture that is more complex than previously assumed. Differential gene expression analysis showed that environmental cues, including temperature, pH, carbon-to-nitrogen ratio and the presence of free fatty acids influence the FASII gene cluster expression in a non-uniform manner. This phenomenon suggests the presence of internal transcription start sites and modular regulation within the cluster. We identified three lipid-responsive transcription factors, MabR_RO_, FadR1_RO_ and FadR2_RO_, that are all capable of binding the *fasII* promoter *in vitro*. DNA binding of FadR1_RO_ and FadR2_RO_ was disrupted by long-chain acyl-CoA molecules, indicating ligand-dependent control. Together, these findings reveal previously unrecognised layers of transcriptional regulation in the *R. opacus* FASII pathway and highlight both conserved and divergent regulatory features within the Mycobacteriales lineage.

**Featured Image:** 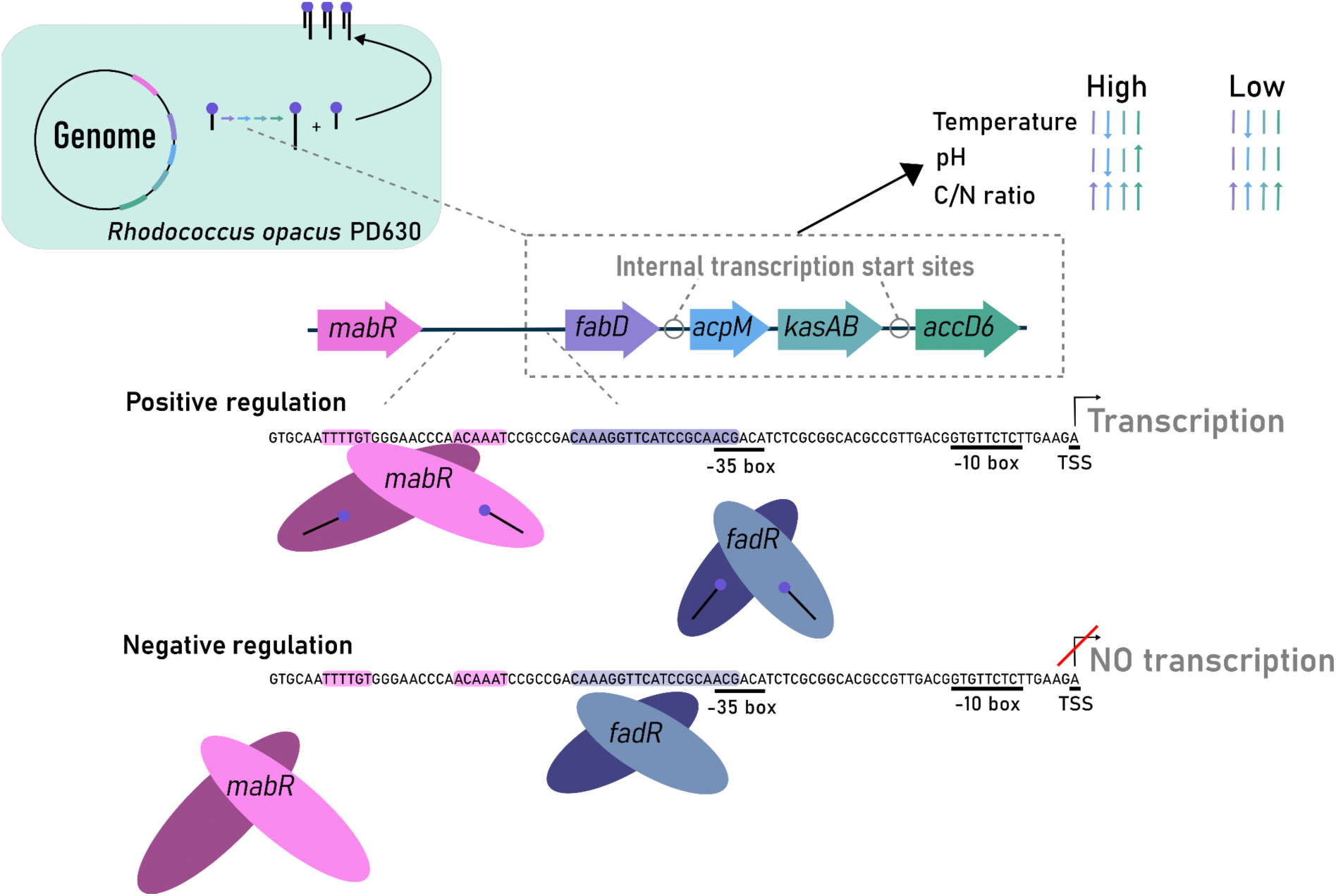

## Introduction

*Rhodococcus opacus* is widely recognised as a model oleaginous bacterium due to its ability to accumulate large amounts of triacylglycerols (TAGs) from diverse and low-cost carbon sources (Alvarez et al., 2016; Anthony et al., 2019; Cappelletti et al., 2020; Donini et al., 2021). Under nitrogen limitation, *R. opacus* PD630 can channel up to 80% of its dry weight into TAGs (Alvarez et al., 2021; Firrincieli et al., 2022), making it an attractive platform for lipid and lipid-derived bioproduction (Alvarez et al., 2021). TAG accumulation is metabolically linked to the *de novo* fatty acid biosynthesis as well as to the production of mycolic acids (Cabruja et al., 2017; Roszkowski et al., 2026), which are very-long-chain α-alkyl-β-hydroxy fatty acids (C_28_ – C_58_) that contribute to the structure of the cell envelope and stress tolerance (Alvarez et al., 2004; Barkan et al., 2009; de Carvalho et al., 2016; Henson et al., 2018; Kellogg James et al., 2001; Stratton et al., 2003; Sutcliffe, 1998). Both fatty acid and mycolic acid synthesis rely on a dual fatty acid synthase (FAS) system in *R. opacus*, consisting of a multifunctional type I fatty acid synthase (FASI), which produces C_16_–C_26_ fatty acyl-CoAs (Holder et al., 2011; Tsai et al., 2017), and a type II system (FASII), a monofunctional enzyme complex encoded by the *kas* (*fabD*-*acpM*-*kasAB*-*accD6*) and *mabA*-*inhA* gene clusters. The latter system elongates long-chain fatty acyl-CoA precursors received from FASI into very long-chain acyl-CoAs (Biswas et al., 2013; Gupta and Singh, 2008; Jakub and Laurent, 2014; Kremer et al., 2002a; Odriozola et al., 1977; Sydor et al., 2013; Takayama et al., 2005; Tsai et al., 2017). Finally, their conversion into mature mycolic acids is mediated by enzymes encoded by the *fadD32*–*pksI*–*accD4* operon (Jain and Ertesvåg, 2022; Marrakchi et al., 2014).

Despite the biotechnological relevance of lipid production in *R. opacus*, regulatory mechanisms governing fatty acid and mycolic acid biosynthesis remain poorly understood. In related mycobacteria, fatty acid synthesis is controlled by multiple transcription factors that regulate *fasII* transcription in response to long-chain acyl-CoAs, including GntR-family FadR (**<u>F</u>**atty **<u>a</u>**cid **<u>d</u>**egradation **<u>r</u>**egulator) (Biswas et al., 2013; Cronan and Subrahmanyam, 1998; Yousuf et al., 2018) and the Mycobacteriales-specific MabR (**<u>M</u>**ycolic **<u>a</u>**cid **<u>b</u>**iosynthesis **<u>r</u>**egulator) (Salzman et al., 2010; Tsai et al., 2017; Zhang and Rock, 2010). Molecular mechanisms employed by FadR and MabR are well-described in *Mycobacterium* spp. Transcriptomic analyses of *M. tuberculosis* have revealed the implication of FadR-type regulators in the coordination of the fatty acid metabolism (Biswas et al., 2013; Ranjan et al., 2015). Five FadR paralogs have been identified, of which only Rv0494 and Rv0586 are known acyl-CoA-responsive regulators that act as a repressor (Biswas et al., 2013; Ranjan et al., 2015; Yousuf et al., 2018). In contrast to FadR, MabR (Rv2242) functions as an activator, as was shown for the ortholog in *Mycobacterium smegmatis* mc^2^155 (Tsai et al., 2017). MabR is essential in *M. tuberculosis* (Salzman et al., 2010) and has been shown to be involved in a larger transcriptional regulatory network: for example, it is proposed to interact with nitrogen-responsive regulators such as NlpR, which also modulates fatty acid biosynthesis (Hernández et al., 2017).

Although *R. opacus* also belongs to the order Mycobacteriales, the presence, function and environmental responsiveness of FadR- and MabR-like regulators remained entirely undocumented in this species, despite these regulators being extensively researched in *E. coli and M. tuberculosis*, respectively (Biswas et al., 2013; Cronan and Subrahmanyam, 1998; Iram and Cronan, 2005; Megalizzi et al., 2024; Ranjan et al., 2015; Salzman et al., 2010; Tsai et al., 2017; van Aalten et al., 2001; Yousuf et al., 2018). Here, we assess the impact of exogenous fatty acid supplementation and environmental stress conditions on transcriptional expression of the *fabD–acpM–kasA/B–accD6* gene cluster, hereafter referred to as the *fasII* gene cluster. In addition, we identify and characterise three MabR- and FadR-like homologous transcriptional regulators in *R. opacus* PD630 — MabR_RO_, FadR1_RO_, and FadR2_RO_ — and determine their putative roles in controlling the mycolic acid biosynthesis pathway. The elucidation of transcription regulatory mechanisms of the *fasII* gene cluster can contribute to future efforts to engineer *R. opacus* as an optimised lipid-producing chassis.

## Materials and Methods

### Bioinformatic analysis

Internal transcription start sites (TSSs) were predicted using RNAseq data from NCBI (GSM1038627 and GSM1038629) processed in RStudio with *GenomicAlignments*, *GenomicFeatures*, *rtracklayer*, *GenomicRanges* and *Rsamtools* (Lawrence et al., 2013, 2009; Morgan et al., 2024; Rstudio Team, 2024). Reads were converted to sorted BAM files and coverage across selected genomic regions was examined to identify positions showing the largest increase in read depth, corresponding to predicted TSSs.

Transcription factor homology searches were performed in the *R. opacus* genome sequence PD630 using pBLAST (Altschul et al., 1990), while orthology searches across selected actinomycetes genomes and comparative functional annotation of differentially expressed genes reported in other studies (Biswas et al., 2013), were conducted using COG resources at the Kyoto Encyclopedia of Genes and Genomes (KEGG) database (Kanehisa et al., 2025).

Protein structures were predicted using AlphaFold2 (Jumper et al., 2021) and AlphaFold3 (Abramson et al., 2024), followed by superimposition using PDB pairwise alignments to obtain Template Modeling (TM) scores (Bittrich et al., 2024). Functional domains were identified with Pfam (Blum et al., 2025) and multiple sequence alignments were generated using the ClustalW tool (Madeira et al., 2024) and visualised in JalView (Waterhouse et al., 2009). Secondary structures were predicted with Psipred (McGuffin et al., 2000). Ligand-binding residues were predicted using IntFold (McGuffin et al., 2019), 3Dligandsite (Wass et al., 2010), BioLIP (Yang, Roy, and Zhang 2013) and COACH (Yang et al., 2013b). DNA-binding residues were predicted using DPbind (Hwang et al., 2007). DNA- and ligand-binding domains were identified with Phyre2 (Kelley et al., 2015). AlphaFold3 was also used to model protein–ligand and protein–DNA interactions, which were further analysed with PDBSum (Laskowski et al., 2018). Binding motif consensus logos were created using WebLogo3 (Crooks et al., 2004).

### Bacterial strains, cultivation and transformation conditions

*Escherchia coli* strain DH5α was utilised for routine cloning and was transformed using an in-house adaptation of the C2987H protocol (New England Biolabs) with a prolonged heat shock time of 45 seconds, revival in Lysogeny Broth (LB) medium (MacWilliams and Liao, 2006) and 2 hours revival time. Cultivation was performed at 37°C and growth was monitored by measuring optical density at 600 nm (OD_600_). Transformant selection was performed on solid LB medium supplemented with 30 µg.mL^-1^ kanamycin or 50 µg.mL^-1^ gentamycin. Heterologous protein expression was performed in *E. coli* strain BL21(DE3) (Studier and Moffatt, 1986).

*R. opacus* PD630 (DSM 44193) (Alvarez et al., 1996) was cultivated in minimal salt medium (MSM) (Saisriyoot et al., 2019) with 10 g.L^-1^ glucose as a carbon and energy source and 2 g.L^-1^ ammonium sulphate as nitrogen source at 29°C and pH 7.0, unless stated otherwise (Alvarez et al., 1996; DeLorenzo and Moon, 2018). To assess the effects of carbon-to-nitrogen (C/N) ratio, pH and temperature, growth was monitored for 48 hours in MSM. For fatty acid exposure experiments, cultures were first grown for approximately 16 hours in MSM under the same carbon and nitrogen conditions. Cultures (20 mL) were then distributed into shake flasks, and 250 µL of a 2 g L⁻¹ C_16_–C_22_ fatty acid stock (dissolved in dimethyl sulfoxide (DMSO)) was added, while 250 µL DMSO was added to the controls. Cultures were incubated for either 3 or 16 hours at 29°C, after which 2 mL of cells was harvested by centrifugation and decantation. All cultures were prepared in triplicate. Pellets were stored at −80°C until they were further processed for RNA extraction.

### RNA extraction and reverse transcriptase quantitative PCR

Cell pellets were resuspended in 600 µL RLT buffer (Qiagen RNeasy kit) and transferred to 2-mL Eppendorf tubes with 300 µL 0.1 mm glass beads. Lysis was performed using a Tissue Lyser II system (2 x 8 minutes, 30 Hz, Qiagen). An equal volume of 70% ethanol was added, and the mixture was processed with an RNeasy RNA extraction kit (Qiagen) according to the manufacturer’s protocol. Residual gDNA was removed using Turbo DNase (Ambion) (Baes et al., 2020). RNA integrity and concentration were assessed by agarose gel electrophoresis (1% bleach) and NanoDrop spectrophotometry, respectively (Aranda et al., 2012). Subsequent cDNA synthesis was preformed from 1 µg RNA with the GoScript Reverse Transcriptase system (Promega).

Reverse transcriptase quantitative PCR (RT-qPCR) was performed in 20-µL reactions containing 8 pmol of each primer, 10 µL GoTaq qPCR Master Mix (Promega) and 1 µL cDNA dilution. A no-template control and a no-RT control were included in each plate. The following program was operated: 3 minutes at 95°C, 40 cycles of 10 seconds at 95°C and 30 seconds at 55°C. RT-qPCR primers (**Supplementary Table S1**) were designed using Primer3 software (Untergasser et al., 2012) and primer efficiency was tested using gDNA as a template (Baes et al., 2020).

RT-qPCR data were analysed using the ΔΔCt method as previously described by Livak & Schmittingen (Livak and Schmittgen, 2001). Differential gene expression of the gene of interest (GOI) was quantified after normalization to the expression of the *ATPClpx* gene (DeLorenzo and Moon, 2018). Technical replicate Ct values were averaged, yielding one log₂ fold change (log₂FC = −ΔΔCt) per biological replicate. Statistical analyses were performed for biological replicates. Mean log₂FC values and 95% confidence intervals (CIs) were calculated using the Student’s *t* distribution. Differential expression was assessed with a one-sample, two-tailed *t*-test comparing log₂FC values to zero, with *p* < 0.05 considered significant. CIs were reported to convey effect size and variability, following recommended practices for quantitative gene expression analysis (Yuan et al., 2006).

### Heterologous protein expression and purification

The *mabR_RO_, fadR1_RO_* and *fadR2_RO_* open reading frames (ORFs) were amplified from *R. opacus* PD630 genomic DNA (gDNA) using gene-specific primers (**Supplementary Table S1**) and cloned into an isopropyl β-D-1-thiogalactopyranoside (IPTG)-inducible pET24a(+) expression vector using a Gibson Assembly approach. The sequence of the plasmid was verified using Sanger sequencing.

FadR1_RO_ and FadR2_RO_ were heterologously expressed as C-terminally His_6_-tagged proteins by inducing with 0.8 mM IPTG at OD_600_ 0.7 and further cultivating at 24 °C for 18 hours. MabR_RO_ was similarly expressed as C-terminal His_6_-tagged recombinant protein by adding 0.1 mM IPTG at OD_600_ 0.7 followed by cultivation at 24 °C for 3 hours. After centrifugation, cell pellets were resuspended in lysis buffer A1 (25 mM Tris, 750 mM NaCl, 5 mM MgCl_2_, 1 mM phenylmethylsulfonyl fluoride (PMSF) and 15 mM imidazole; pH 7.4) or A2 (50 mM Tris, 750 mM NaCl, 1 mM PMSF and 15 mM imidazole; pH 7.4) for MabR_RO_ and FadR1_RO_/FadR2_RO_, respectively. After sonication for 15 minutes at 20% amplitude (Vibracell 75043, Bioblock Scientific), the supernatant was subjected to purification using Ni-NTA (HisTrap FF 1mL, VWR) affinity chromatography on the ÄKTA purification system (Cytiva). Elution was performed with buffer B1 (25 mM Tris, 750 mM NaCl, 5 mM MgCl_2_, 1 mM PMSF and 750 mM imidazole; pH 7.4) or B2 (50 mM Tris, 750 mM NaCl, 1 mM PMSF and 750 mM imidazole; pH 7.4) for MabR_RO_ and FadR1_RO_/FadR2_RO_, respectively. Different peak fractions were collected, analysed using sodium dodecyl sulfate polyacrylamide gel electrophoresis (SDS-PAGE) and stored in buffer C (50 mM Tris, 5% (v/v) glycerol). The final yield amounted ∼0.4 mg or ∼8 mg per 300 mL bacterial culture for MabR_RO_ and FadR1_RO_/FadR2_RO_, respectively.

### Electrophoretic mobility shift assays

Electrophoretic mobility shift assays (EMSAs) were performed to analyse interactions between the purified proteins and a 292-bp probe representing the native *fasII* promoter region. This probe was PCR-amplified from *R. opacus* PD630 gDNA using primers PL_0048 and PL_0049 (**Supplementary Table S1**), of which one was end-labelled with [γ-^32^P]-ATP (3000 Ci mmol^-1^) with T4 polynucleotide kinase. The ^32^P-labelled DNA probe was gel-purified and combined with protein in 10 µL binding buffer D1 (25 mM Tris/HCl pH 8, 1 mM PMSF, 5% (v/v) glycerol, 5 mM MgCl_2_, 150 mM NaCl) (Salzman et al., 2010) or D2 (10 mM Tris/HCl pH 8, 0.5 µg non-specific DNA mL^-1^, 5% (v/v) glycerol, 1 mM EDTA, 10 mM NaCl) (Ranjan et al., 2015) for MabR_RO_ and FadR1_RO_/FadR2_RO_, respectively. Reaction mixtures were incubated at 28°C for 60 minutes or at 30°C for 30 minutes for MabR_RO_ and FadR1_RO_/FadR2_RO_, respectively. To assess ligand effects, 45 µM arachidoyl-CoA was added to the reaction mixture. DNA-protein complexes were resolved on a 6% (w/v) non-denaturing polyacrylamide gel in 0.2X TBE buffer at 110 V at 4°C and visualised using a Phosphorimager (BioRad).

## Results

### The fasII gene cluster exhibits a non-operonic architecture and limited transcriptional responsiveness to extracellular fatty acids

To investigate the transcriptional architecture of the *fasII* gene cluster in *R. opacus* PD630, TSSs were predicted *in silico* based on two available RNA-seq datasets (**Figure 1A**). Interestingly, a gene encoding a putative MabR homolog (*mabR*) is located adjacently upstream to the *fasII* gene cluster. Individual TSSs were predicted for *mabR*, *fabD*, *acpM* and *accD6* promoter regions, consistent with the relative long intergenic regions preceding these genes (**Figure 1A**). For *acpM*, two alternative TSSs were identified, separated by 93 bp. In contrast, no TSS was predicted for the *kasAB* gene, indicating that it is co-transcribed with *acpM*. These observations suggest that the *fasII* gene cluster is not transcribed as a single operonic unit, which is supported by the marked differences in RNA-seq read coverage for *acpM*, *kasAB* and *accD6* as compared to *mabR* and *fabD* (**Figure 1A**).

**Figure 1.**
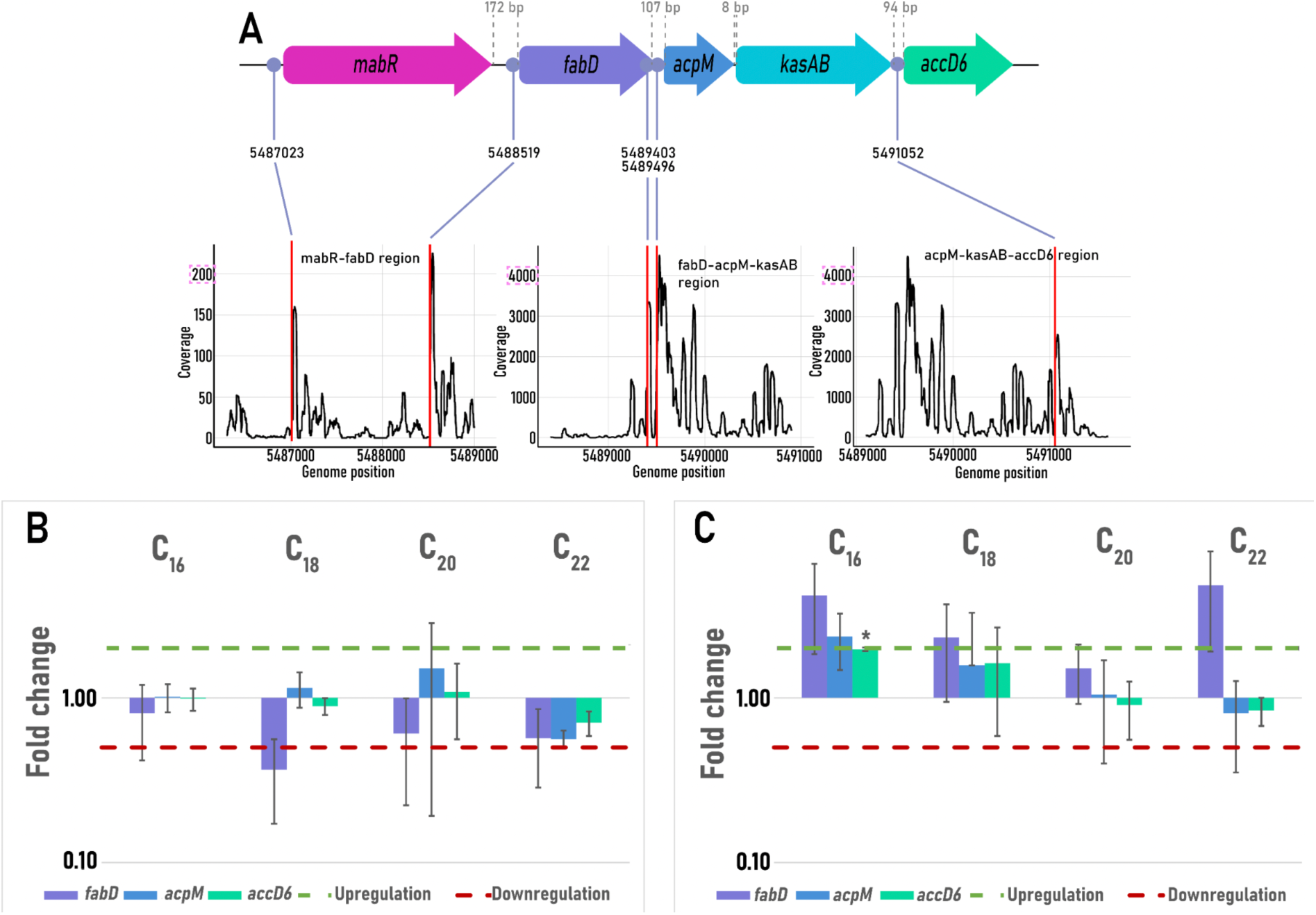
Transcriptional structure and fatty-acid responsiveness of the *R. opacus fasII* gene cluster. (**A**) Prediction of transcription start sites for the *fasII* gene cluster and the *mabR_RO_* gene. Gene abbreviations in the gene cluster: malonyl-CoA:AcpM transacylase (*fabD*), mycobacterial acyl carrier protein (*acpM*), β-ketoacyl-AcpM reductase (*kasAB*) and acetyl-CoA carboxylase (*accD6*). The predicted transcription start site coordinates are presented alongside coverage plots for various genomic regions. To emphasise the large differences in coverage scale between regions, the maximum annotated read coverage is indicated by a pink dashed line. (**B & C**) Relative transcriptional expression after supplementation with 100 mg.L^-1^ fatty acids for 3 (**B**) and 16 hours (**C**) of induction. Significant differences in expression compared to the control group (without fatty acid supplementation) at α = 0.05, as determined by a Student’s T-test, are indicated with an asterisk (*). Number of biological and technical replicates were three, except for the C_16_ condition, for which two biological replicates were included. Error bars indicate standard deviation.

In light of the functional role of the FASII system, we hypothesise that transcriptional expression of the *fasII* gene cluster is responsive to the presence of free long-chain fatty acids in the culture medium as these are taken up by the cell and contribute to the long-chain fatty acyl-CoA or ACP pool (Arora et al., 2009). To test this, relative transcriptional expression of the *fabD*, *acpM* and *accD6* genes was tested in absence and presence of C_16_, C_18_, C_20_ or C_22_ saturated fatty acids with a RT-qPCR approach (**Figure 1B-C**). At 16 hours after fatty acid supplementation (**Figure 1C**), a modest increase in transcriptional expression of *fabD*, *acpM* and *accD6* was observed in case of C_16_ fatty acids, albeit only significant for the latter gene. In contrast, supplementation with longer-chain fatty acids (C_18_, C_20_ or C_22_) resulted in little or no transcriptional response. Moreover, a 3-hour induction experiment revealed an opposite trend, with generally unchanged or slightly reduced expression levels following fatty acid supplementation (**Figure 1B**). These observations do not support the initial hypothesis that transcriptional expression of the *fasII* gene cluster is induced in response to extracellular long-chain fatty acids under the tested conditions.

### Environmental stress responses support a non-canonical transcriptional organisation of the fasII cluster

To further elucidate transcriptional regulation of the *fasII* gene cluster, the effects of pH, temperature, and carbon/nitrogen (C/N) ratio on expression of *fasII* cluster genes were investigated by RT-qPCR (**Figure 2, Supplementary Table S2**). Specifically, pH levels of 5.5 or 8, temperatures of 22°C or 34°C and C/N ratios of 38 or 184 were compared to the optimal growth conditions at pH 7 (Saisriyoot et al., 2019), 29°C (Alvarez et al., 1996; DeLorenzo and Moon, 2018) and a C/N ratio of 9.5 (**Supplementary Figure S2**).

**Figure 2.**
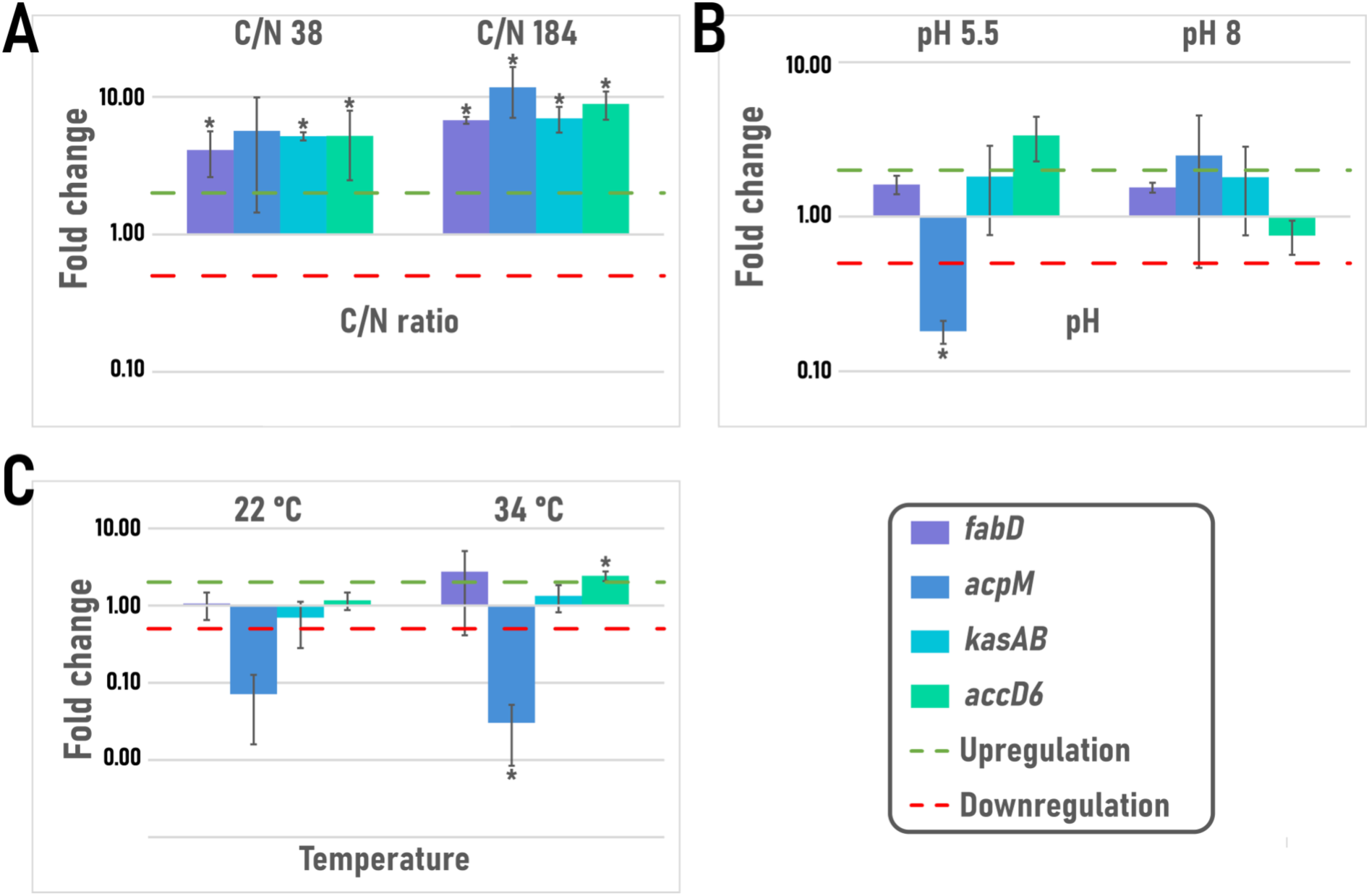
Relative transcriptional expression of *fasII* cluster genes at different environmental perturbations. (**A**) Effect of different C/N ratios relative to the control C/N ratio of 9.5. (**B**) Effect of pH relative to the control pH of 7. (**C**) Effect of temperature relative to the control temperature of 29°C. Significant differences in expression compared to the respective control conditions at α = 0.05, as determined by a Student’s T-test, are indicated with an asterisk (*). Biological and technical replicates were three for all conditions except for the pH experiments, for which only two biological replicates were included. Error bars indicate standard deviation.

Increased C/N ratios led to consistent upregulation of all genes within the *fasII* cluster, with an overall stronger induction observed at a C/N ratio of 184 than at 38 (**Figure 2A**). A slightly alkaline pH of 8, above the optimal growth pH of 7, did not significantly affect *fasII* transcriptional expression. In contrast, exposure to acidic conditions (pH 5.5) resulted in a significant downregulation of *acpM* expression, while expression levels of *fabD*, *kasAB* and *accD6* remained largely unaffected (**Figure 2B**). Similarly, temperature shifts relative to the optimal growth temperature of 29°C primarily affected *acpM* expression, while the remaining *fasII* genes showed limited transcriptional responses (**Figure 2C**). At suboptimal temperature (22°C), *acpM* expression was significantly reduced, whereas at 34°C downregulation of *acpM* was accompanied by a modest but significant downregulation of *accD6*.

### Prediction of putative transcription regulators of the fasII gene cluster

To identify candidate transcription regulators that could mediate long-chain acyl-CoA-dependent control of the *fasII* gene cluster, we searched for homologs of the known mycobacterial GntR-family FadR-type regulators Rv0586 and Rv0494. Bioinformatic analysis identified 18 GntR family proteins in *Rhodococcus opacus* PD630 belonging to the FadR subfamily. Among these, two were predicted to be candidate acyl-CoA-responsive regulators based on their sequence similarity to Rv0586 and Rv0494: FadR1_RO_ (geneID: Pd630_LPD06939) and FadR2_RO_ (Pd630_LPD05140) (**Figure 3, Supplementary Figure S3**). FadR2_RO_ was the closest homolog of FadR_Ecoli_, although overall sequence identity was low (22.9% sequence identity). Structural comparison showed that FadR2_RO_ is most similar to Rv0586, followed by Rv0494, whereas FadR1_RO_ showed lower similarity to the mycobacterial paralogs (**Figure 3A**).

**Figure 3.**
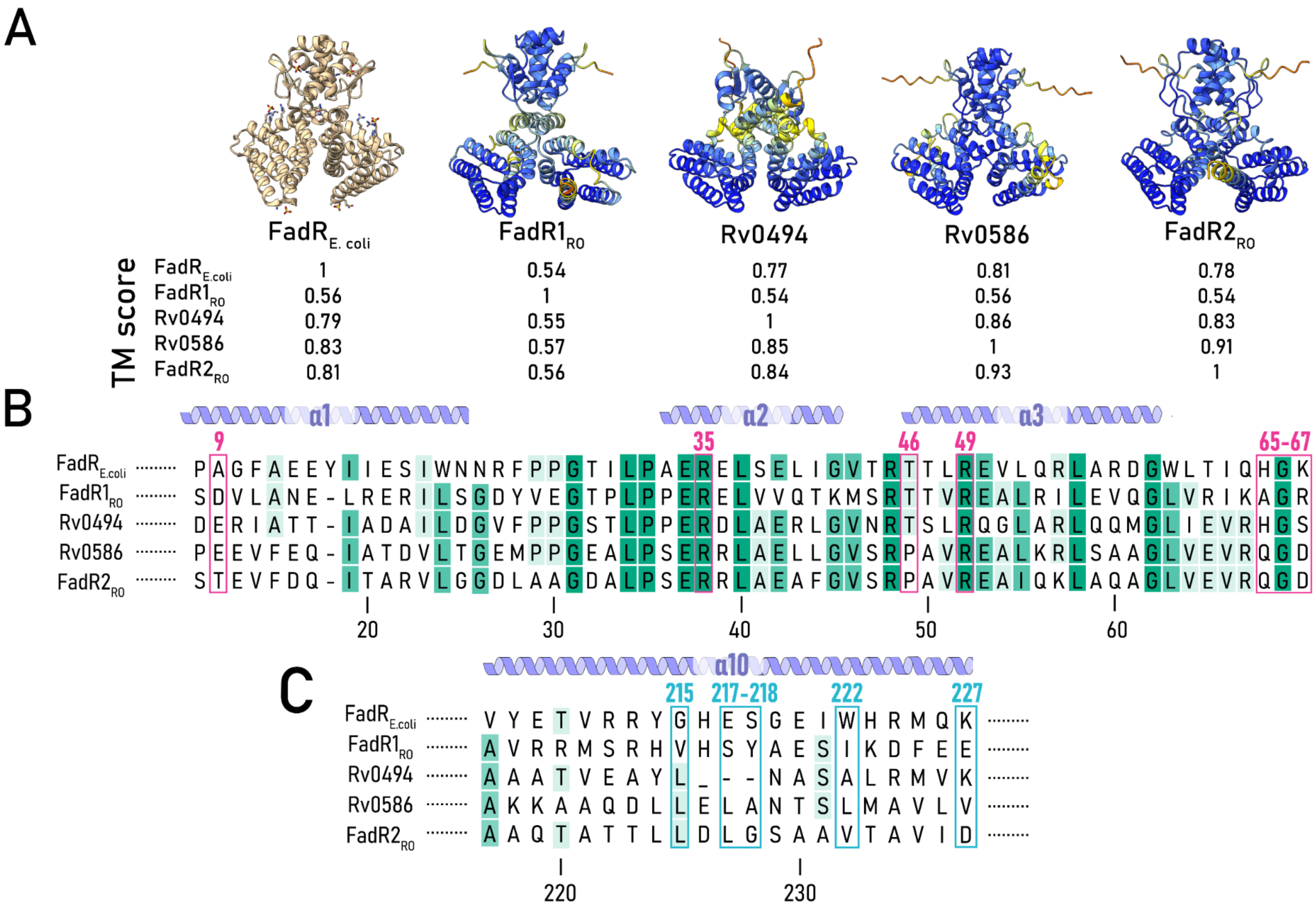
Structural and sequence comparison of FadR homologs from *R. opacus*, *M. tuberculosis* and *E. coli*. (**A**) Predicted structures of FadR1_RO_ and FadR2_RO_ from *R. opacus* PD630, Rv0586 and Rv0494 from *M. tuberculosis*, compared with the crystal structure of FadR_Ecoli_ (1H9T) (van Aalten et al., 2001). Pairwise TM scores are shown below the structures, with values ranging from 0 to 1, with higher values indicating greater structural similarity. Because TM scores are normalised relative to the selected reference structure and length, reciprocal comparisons may yield slightly different values. (**B**) Multiple sequence alignment of the N-terminal domain of the FadR homologs. Alpha helices, based on the FadR_Ecoli_ structure are displayed above the alignment. Position numbering is with respect to FadR_Ecoli_. DNA-binding residues are highlighted in magenta. (**C**) Multiple sequence alignment of the C-terminal domain of the different homologs. Ligand-binding residues are highlighted in blue.

Multiple sequence alignment revealed a high level of conservation of residues demonstrated to be important for DNA binding in FadR_Ecoli_ within the N-terminal helix-turn-helix (HTH) domain (Ala9, Arg35, Thre46, Arg49, His65, Gly66 and Lys67) (Biswas et al., 2013; Iram and Cronan, 2005; Raman et al., 1997; van Aalten et al., 2001; Yousuf et al., 2018) (**Figure 3B**). Indeed, on several of the corresponding positions in the actinobacterial homologs, conserved or similar residues are present, supporting a shared DNA-binding function. In contrast, residues in the C-terminal ligand-binding region of FadR_Ecoli_ (Raman et al., 1997) were found to be poorly conserved across the homologs (**Figure 3C**), with only limited conservation around Leu215. This divergence is consistent with potential differences in ligand-binding affinity or specificity among FadR homologs (Iram and Cronan, 2005).

In addition to the FadR homologs, a MabR homolog, MabR_RO_, was predicted to be encoded divergently with respect to the *fasII* gene cluster (**Figure 1A**). It was found to display 63% sequence identity with the PucR-family MabR homolog in *M. tuberculosis*, supported by the prediction of a typical N-terminal GGDEF domain and C-terminal HTH motif, involved in gene regulation and DNA binding, respectively (Lu et al., 2020; Ryjenkov et al., 2005). MabR homologs were found to be well-conserved across the Mycolata taxon, with an overall higher degree of sequence conservation for the C-terminal HTH motif as compared to the N-terminal domain (**Figure 4A-B**). Indeed, all Mycolata MabR homologs harbor a stretch of residues, [353-E][362-HPNTVRYRLK-371] (**Figure 4A**), which are confirmed to be DNA-binding determinants by a structural model and protein-DNA interaction predictions (**Supplementary Dataset S2, Supplementary Figure S4 & S5**). Similarly to the *M. tuberculosis* MabR regulator (Tsai et al., 2017), MabR_RO_ was predicted to interact with long-chain acyl-CoA molecules in its N-terminal domain via hydrogen bonds and other non-covalent bonds (**Figure 4B, Supplementary Figure S5, Supplementary Dataset S1**). As *M. tuberculosis* MabR can adopt both dimeric and tetrameric states, with DNA binding promoting a shift towards the dimeric form, MabR_RO_ was modelled as a dimer and tetramer, with and without ligands (DNA and/or a long-chain acyl-CoA ligand) (**Figure 4C, Supplementary Figure S5**). The predicted MabR_RO_ dimer adopts an extended, relatively planar conformation. According to the structural interaction analysis (**Supplementary Figure S5 & S6**), the dimer interface involves 26 residues, forming 4 salt bridges, 6 hydrogen bonds and 107 non-bonded interactions. The tetrameric model consists of two dimers arranged symmetrically along their longest axis. The predicted dimer-dimer interface contains 6 salt bridges, 9 hydrogen bonds and approximately 258–265 non-bonded interactions, consistent with an extensive stabilizing interface between the two dimers.

**Figure 4.**
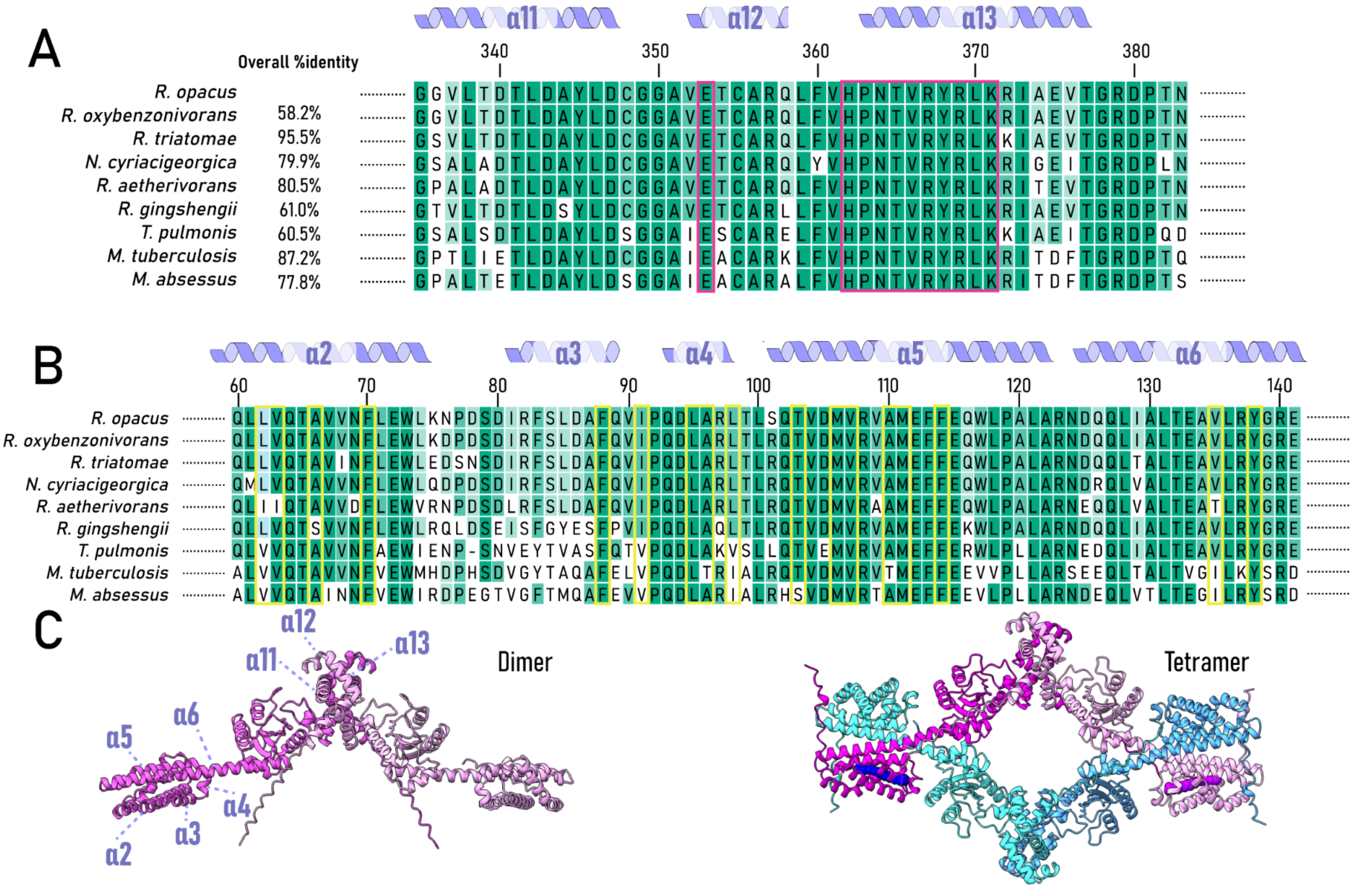
Sequence comparison of MabR homologs from different members of the Mycolata taxon and structural modelling of MabR_RO_. (**A**) Multiple sequence alignment of the C-terminal domains of MabR homologs. Alpha helices are based on the prediction for MabR_RO_ and are indicated above the alignment. Position numbering is with respect to MabR_RO_. Predicted DNA-binding residues are highlighted with a magenta box. (**B**) Multiple sequence alignment of the N-terminal domain of MabR homologs. Alpha helices are based on the prediction for MabR_RO_ and are indicated above the alignment. (**C**) Structure prediction of MabR_RO_ as dimer and tetramer with an acyl-CoA ligand, indicated in a space-filling depiction.

### Prediction of FadR and MabR binding motifs in the fasII promoter region

We next employed a bioinformatic approach to predict putative binding motifs for the identified transcription regulators. With regards to FadR-like regulators, a FadR binding motif search was performed for selected sequences from the *R. opacus* PD630 genome sequence, focused on upstream regions (< 350bp) of genes homologous to FadR regulon genes in *M. tuberculosis* (Biswas et al., 2013) and employing the consensus binding motif of FadR*_Ecoli_* (Gao et al., 2017) as a query. The promoter regions of *fadR1_RO_* and *fadR2_RO_* were included as well, given that GntR-type FadR regulators are known to perform autoregulation. The majority of selected genes showed the presence of one or multiple potential FadR-binding sites in their upstream regions with the following consensus binding motif: N<u>A</u>cN<u>GGT</u>NNNN<u>CC</u>NNNN (**Figure 5A, Supplementary Table S3**). For MabR, we started from the previously described palindromic binding motif in *M. tuberculosis* T<u>TTTGT</u>(N)_9_<u>ACAAA</u>**T** (Salzman et al., 2010) and performed a genome-wide search for this motif in the *R. opacus* genome, leading to the identification of eight potential MabR binding sites upstream of genes encoding membrane proteins, enzymes and transcription factors, in addition to the *fasII* gene cluster (**Figure 5B, Supplementary Table S4**), suggesting a broader regulatory role.

**Figure 5.**
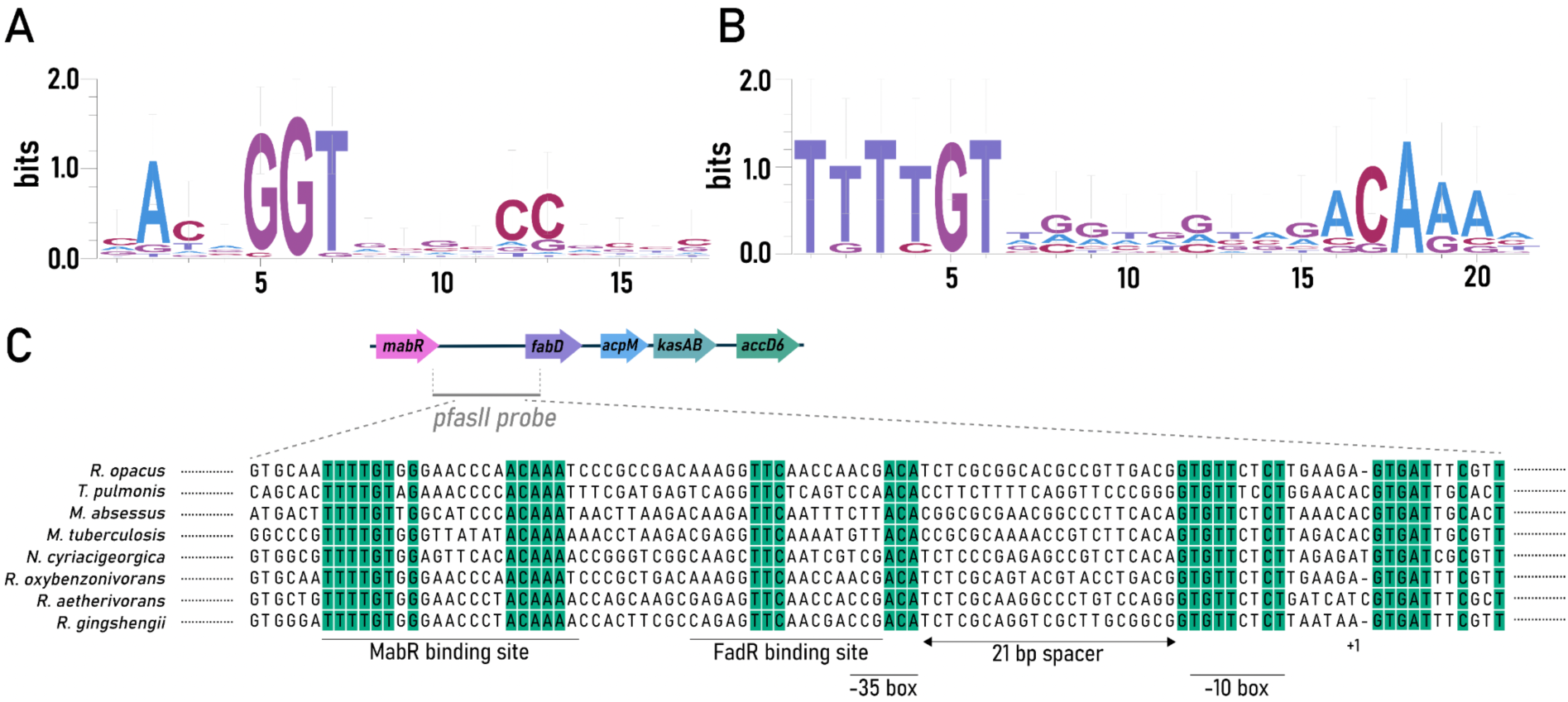
Predicted FadR and MabR binding motifs and conservation of the *mabR*–*fabD* intergenic region. (**A**) Sequence logo representing the predicted FadR binding motif, generated from putative FadR binding sites identified in selected upstream regions of the *R. opacus* PD630 genome (**Supplementary Table S3**). Nucleotide frequencies at each position are visualised to highlight conserved bases and motif symmetry. **(B**) Sequence logo representing the MabR binding motif derived from experimentally validated and predicted MabR binding sites in the *R. opacus* PD630 genome (**Supplementary Table S4**), illustrating conserved sequence features of the predicted MabR-binding motif. (**C**) Multiple sequence alignment of the intergenic region between *mabR* and *fabD* across representative species within the order *Mycobacteriales*. Conserved nucleotides shared among the analysed species are highlighted in green, indicating putative regulatory elements and conserved transcription factor binding sites within this locus. A 21-bp spacer separates the predicted −35 and −10 promoter elements in the aligned *fabD* promoter region.

Based on the conserved regulation of homologous *fasII* gene clusters in other Mycolata species, we hypothesised that FadR1_RO_, FadR2_RO_ and MabR may target the *fabD* promoter region in *R. opacus*. Therefore, we set out to predict putative binding sites for the respective regulators in this promoter region using the identified consensus sequences as well as a phylogenetic footprinting approach (**Figure 5C**). Putative -35 and -10 promoter elements could be inferred in the *fabD* promoter region, showing good agreement with the established *Rhodococcus* consensus promoter motifs (TTGNNN for the −35 box and (T/C)GNNA(A/C)AAT for the −10 box) (Jiao et al., 2018). The predicted MabR binding site was located 31 bp upstream of the -35 box. In addition, a single conserved FadR binding site was identified, located closer to the promoter and even overlapping 3 bp with the -35 box.

### MabR_RO_, FadR1_RO_ and FadR2_RO_ interact with the fabD promoter region in vitro

*In vitro* binding of MabR_RO_, FadR1_RO_ and FadR2_RO_ to the *fasII* promoter region was investigated by EMSA using purified proteins and a 292-bp ^32^P-labelled DNA fragment representing the *fabD* promoter region from 172 bp upstream to 119 bp downstream of the TSS, covering the entire *mabR-fabD* intergenic region (**Figure 6**). The influence of arachidoyl-CoA on protein-DNA complex formation was also investigated, because long-chain acyl-CoA molecules are known ligands of FadR- and MabR-type transcription regulators (Ranjan et al., 2015).

**Figure 6.**
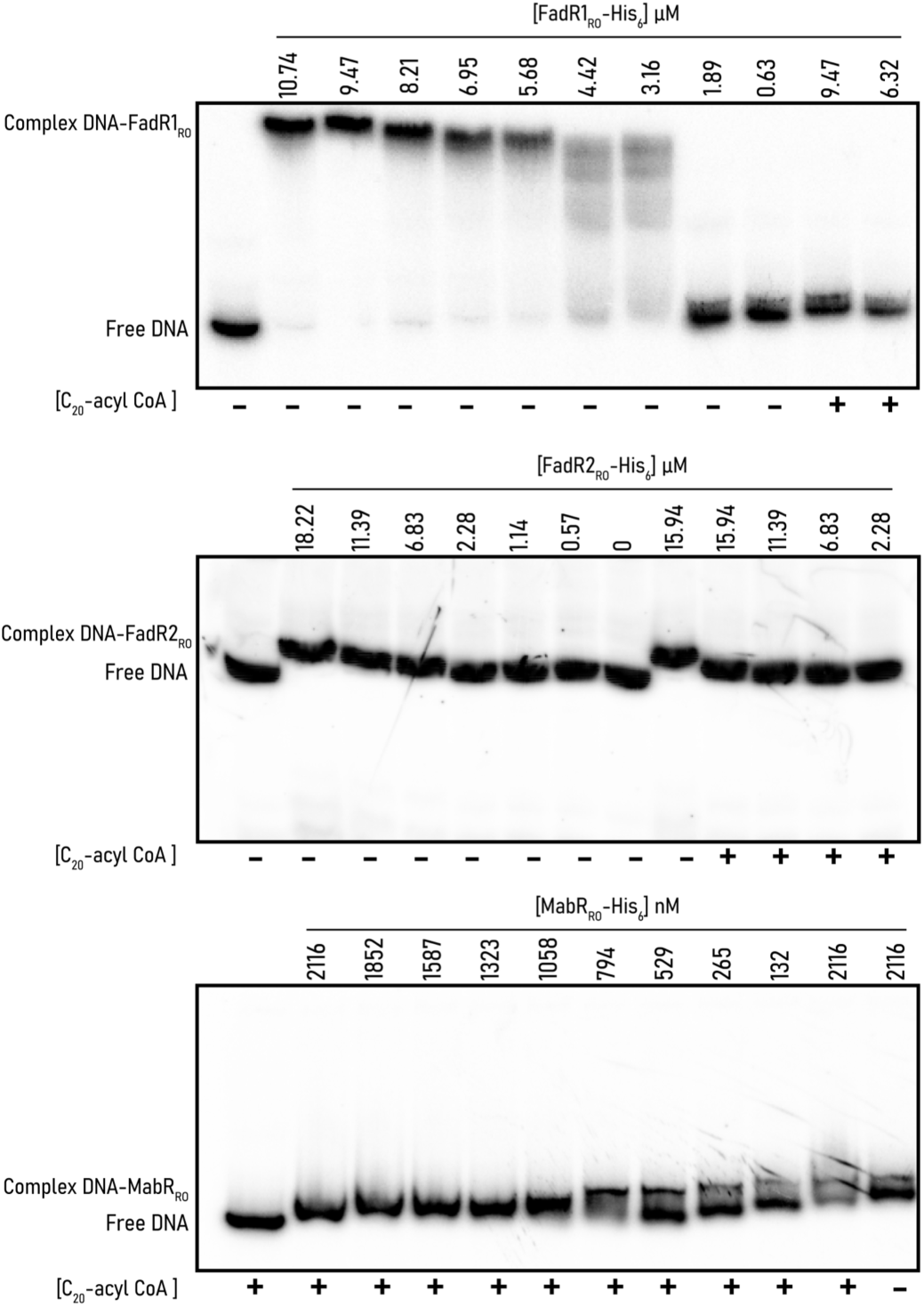
Interaction of FadR1_RO_, FadR2_RO_ and MabR_RO_ with the *fasII* promoter region. EMSAs were performed using purified recombinant FadR1_RO_, FadR2_RO_ and MabR_RO_ and a ^32^P-labelled DNA fragment covering the *mabR*-*fabD* intergenic region. Protein concentrations are indicated above each lane. Arachidoyl-CoA (C_20_-acyl-CoA) was added, as indicated with a “+” symbol below each lane. The positions of protein-DNA complexes were assigned based on their reduced electrophoretic mobility relative to the free DNA probe.

EMSAs demonstrated that recombinant FadR1_RO_ and FadR2_RO_ are both capable of interacting with the *fasII* DNA probe in a concentration-dependent manner (**Figure 6**). Although complex formation was observed only at estimated μM protein concentrations, which is relatively high for a sequence-specific transcription regulator, this may reflect partial loss of protein activity during purification or suboptimal *in vitro* binding conditions. Notably, the FadR1_RO_-DNA and FadR2_RO_-DNA complexes displayed markedly different electrophoretic mobilities: FadR1_RO_ caused a pronounced shift, whereas FadR2_RO_ caused only a small reduction in probe migration velocity (**Figure 6**). This difference suggests that the two regulators may form complexes with different stoichiometries or conformations, despite their predicted structural similarity (**Figure 3**). In both cases, addition of arachidoyl-CoA disrupted formation of the FadR1_RO_-DNA and FadR2_RO_-DNA complexes, indicating acyl-CoA-dependent modulation of DNA binding (Ranjan *et al*. 2015)

MabR_RO_ also interacted with the *fasII* DNA probe, and although the interaction was also observed in the nM range, the observed shift was modest compared to that caused by FadR1_RO_ (**Figure 6**). Under the tested conditions, complex formation appeared more pronounced in the presence of arachidoyl-CoA, suggesting that this long-chain acyl-CoA may stabilise MabR_RO_ binding rather than disrupting it (Salzman *et al*. 2010).

## Discussion

Our results show that environmental perturbations affect *fasII* gene expression in *R. opacus* PD630 in a condition- and gene-specific manner. Increased C/N ratios resulted in coordinated upregulation of the tested *fasII* genes, suggesting that carbon-rich conditions may increase the demand for fatty acid and mycolic acid biosynthesis. By contrast, pH and temperature shifts mainly affected *acpM* and, to a lesser extent, *accD6*, indicating that individual genes within the cluster do not respond uniformly to environmental stress. Such responsiveness has also been observed in other Mycobacteriales. In *M. tuberculosis*, temperature changes affect mycolic acid content and composition (Kellogg et al. 2001, Alvarez et al. 2004, Barkan et al. 2009) and are accompanied by elevated FASII activity (Baba et al., 1989; Kremer et al., 2002), while acidic pH has been shown to modulate expression of fatty acid and mycolic acid genes in an exposure-dependent manner (Fisher et al., 2002). Together, these observations suggest that environmental modulation of fatty acid and mycolic acid biosynthesis is a broader adaptive feature within Mycobacteriales, although the specific transcriptional responses differ between species and conditions (Alvarez et al., 1996; Baker et al., 2014; Fisher et al., 2002; Hernández et al., 2017; Janßen et al., 2013; Rohde et al., 2007).

A modular transcriptional organisation of the *fasII* gene cluster is consistent with predicted internal TSSs upstream of *acpM* and *accD6*. Also in *M. tuberculosis* and *M. smegmatis*, *accD6* is controlled by its own promoter, with altered *accD6* expression influencing growth, mycolic acid levels and morphology (Liu et al., 2018; Pawelczyk et al., 2011). Such layered transcriptional complexity highlights the need for fine-tuned control of fatty acid and mycolic acid biosynthesis, consistent with the central role of mycolic acids in stress tolerance across Mycobacteriales (Alvarez et al., 2004; Barkan et al., 2009; Kellogg et al., 2001).

Despite the more complex transcriptional architecture, the divergent *mabR*-*fabD* intergenic region remains a plausible regulatory “hotspot” for transcriptional control. Also in homologous mycobacterial systems, *fasII*-associated genes are controlled by different acyl-CoA-responsive transcription regulators. In *R. opacus* PD630, we identified three candidate regulators with potential relevance, one MabR homolog and two FadR homologs, and demonstrated that all interact with the *mabR*-*fabD* intergenic region. Although the precise binding site locations occupied by each regulator remain to be experimentally defined, predicted binding motifs are located at positions compatible with an activator and repressor role for MabR and FadR, respectively. Indeed, the predicted MabR binding site lies upstream of the -35 promoter element, in line with a class I activator function as described for MabR in *M. tuberculosis* (Bervoets and Charlier, 2019; Tsai et al., 2017). The observation that two FadR paralogs interact with the same promoter region is particularly interesting, because it suggests an additional layer of complexity not previously demonstrated for this locus in *Rhodococcus*. Whether these two paralogs compete for the same binding site, *i.e.* the conserved promoter-overlapping FadR binding motif, or act under different physiological conditions, remains unresolved.

EMSA analysis further indicated that all three candidate regulators respond to long-chain acyl-CoA, although in different ways. Arachidoyl-CoA disrupted formation of the FadR1_RO_-DNA and FadR2_RO_-DNA complexes, consistent with the established mechanism of FadR-type regulators in which ligand binding reduces DNA-binding activity and relieves repression (Ranjan et al., 2015). In contrast, MabR_RO_-DNA complex formation appeared more pronounced in the presence of arachidoyl-CoA, suggesting that long-chain acyl-CoA may stabilise MabR_RO_ binding to the *mabR*-*fabD* intergenic region, in line with the ligand-responsive role described for MabR in mycobacteria (Megalizzi et al., 2024; Tsai et al., 2017). This opposite response of FadR- and MabR-type regulators provides a regulatory logic in which intracellular long-chain acyl-CoA levels could modulate transcription of the *fasII* gene cluster through both positive and negative regulatory inputs. The absence of a clear transcriptional induction of the *fasII* genes upon supplementation with extracellular fatty acids does not necessarily contradict this model, as the tested conditions may not have resulted in sufficient or sustained changes in the intracellular acyl-CoA pool, influenced by fatty acid uptake, activation into acyl-CoA and metabolic flux through competing lipid pathways. In conclusion, the findings in this study support a regulatory model in which *R. opacus* fine-tunes mycolic acid production by integrating intracellular acyl-CoA availability with broader metabolic and environmental cues.

## Acknowledgments

The authors would like to thank Rani Baes for helping with the RT-qPCR data analysis.

## Data availability statement

All data supporting the findings of this study are available in the Supplementary Materials and in Zenodo at 10.5281/zenodo.20538677.

