## Supplementary Materials for "Transcriptional regulation of the type II fatty acid synthase complex-encoding gene cluster in *Rhodococcus opacus*"

Leemans *et al.*

**Supplementary Materials**

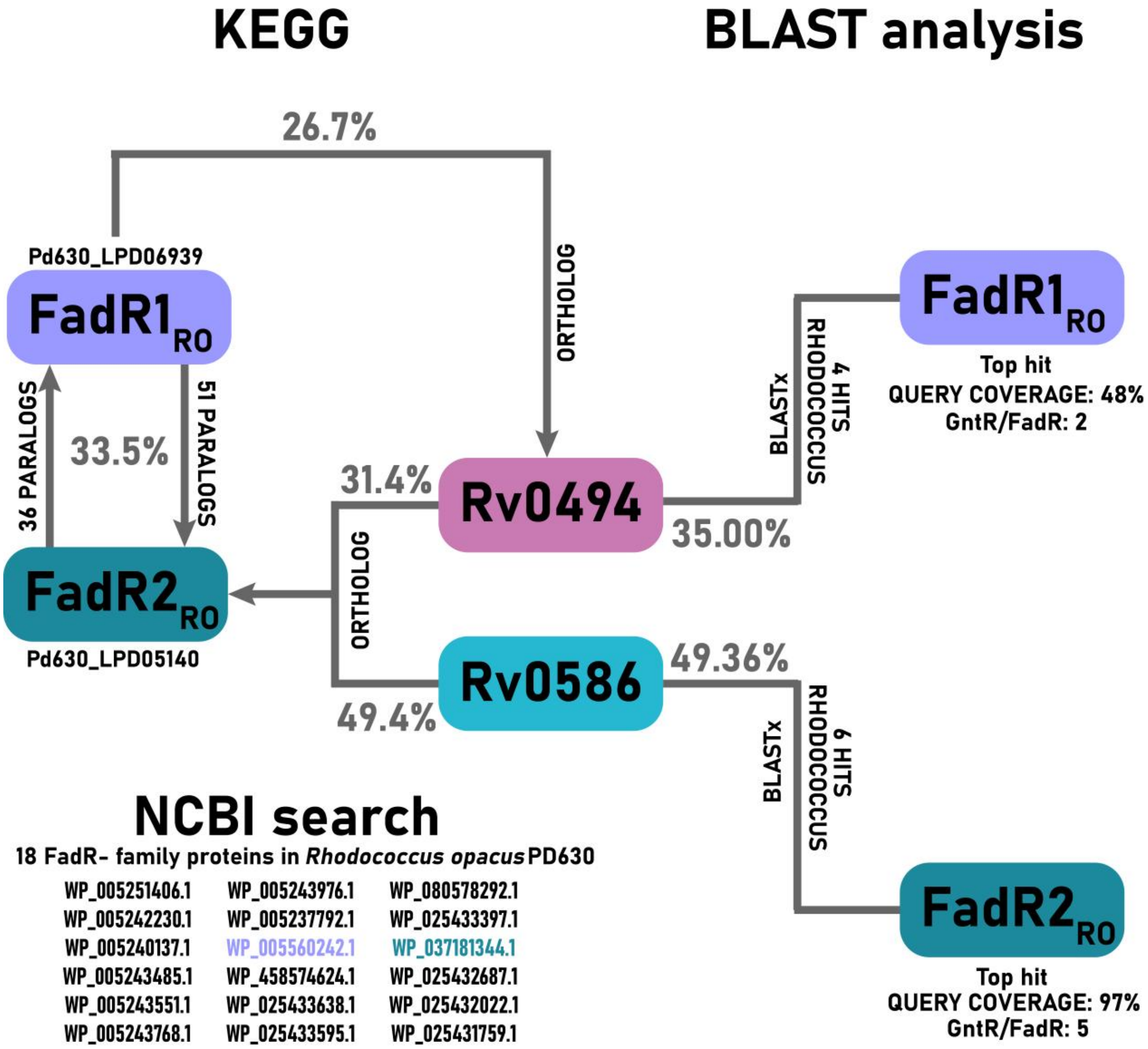


**Supplementary Figure S1.** BLAST and KEGG-based identification of FadR-like proteins in *Rhodococcus opacus* PD630. KEGG orthology and BLAST analyses identified two candidate FadR-type regulators in *R. opacus* PD630, LPD05140 ($FadR2_{RO}$) and LPD06939 ($FadR1_{RO}$), showing sequence identities of ~26–50% to Rv0494 and Rv0586. BLASTx results indicate higher query coverage and similarity for FadR2 compared to FadR1. Both proteins are associated with GntR/FadR regulator groups and are part of larger paralogous sets (36–51 members). NCBI database searches identified 18 FadR-family proteins in the PD630 genome, listed at the bottom of the figure.

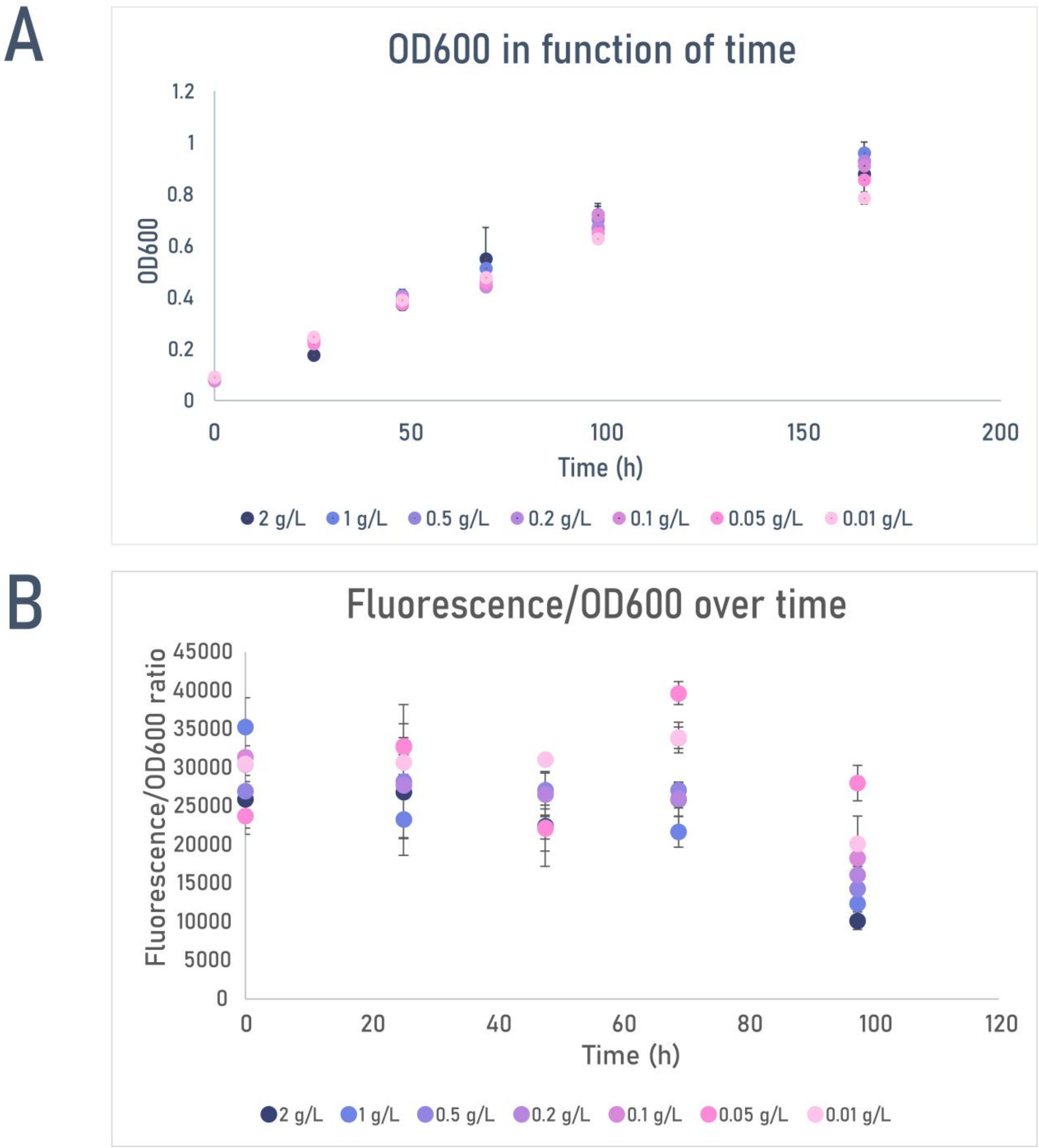


**Supplementary Figure S2**. Determination of the optimal C/N ratio for growth and lipid production in *R. opacus* PD630. (**A**) Growth ($OD_{600}$) over time at different ammonium sulphate concentrations, corresponding to C/N ratios of 9.6 (1 g/L), 38 (0.2 g/L), and 184 (0.05 g/L). (**B**) Lipid accumulation assessed as Nile Red fluorescence normalized to $OD_{600}$ at varying nitrogen levels. Fluorescence was measured using a Synergy HTX plate reader ($\lambda_{ex}$ 485 nm; $\lambda_{em}$ 575 nm).

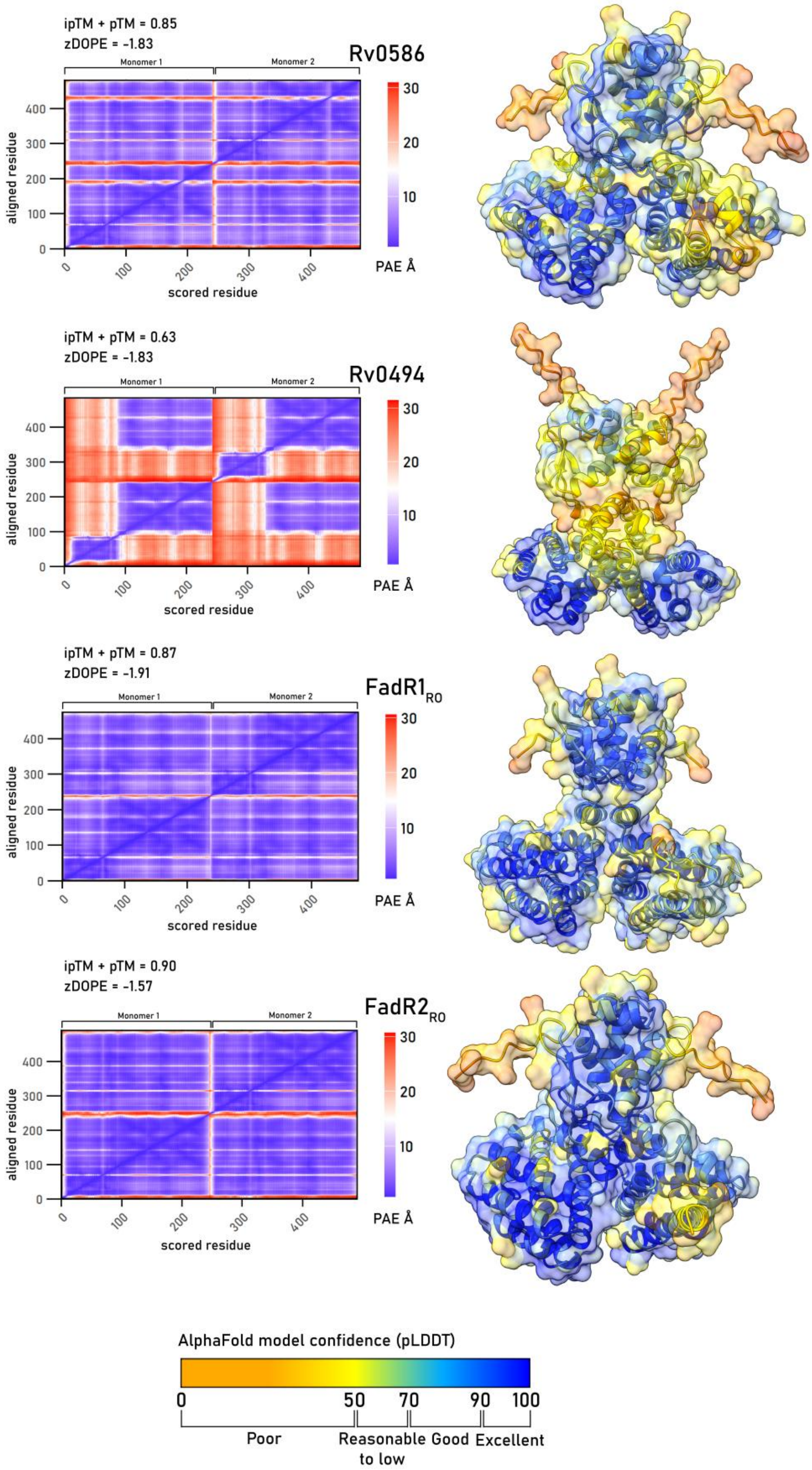


**Supplementary Figure S3**. Legend on next page.

**Supplementary Figure S3**. Cartoon representations of the models generated by AlphaFold3 (Rv0586, Rv0494, FadR1$_{RO}$, FadR2$_{RO}$) in dimeric and tetrameric formation, with or without ligand. The predictions are coloured according to the predicted local distance difference test (pLDDT), which indicates the local quality of the model, as shown in the legend. For each structure, the predicted aligned error (PAE), normalised discrete optimised protein energy (zDOPE), and overall predicted template modelling (pTM) scores are displayed. The PAE predicts the positional error for each residue x (scored residue) in a protein structure when aligned with its corresponding residue y (aligned residue) in the true structure. A zDOPE score below -1 suggests that the atom pair distances in the model are similar to those in a large sample of known protein structures, with at least 80% of the model's Cα atoms within 3.5 Å of their correct positions. As all the predicted structure are multimers, the pDockQ and AlphaFold-Multimer model confidence (0.8ipTM + 0.2pTM). The pTM score (ranging from 0 to 1) is the predicted TM score for a superposition between the predicted and the hypothetical true structure, indicating the accuracy of the prediction within a single chain. Contrary to the interface pTM (ipTM, ranging from 0 to 1), that evaluates the similarity between the true and predicted structures over interfacing residues, reflecting the accuracy of the prediction for a complex. The pDockQ score (ranging from 0 to 1) is another confidence criterion for protein complexes, considering the number of interfacing residues and their pLDDT score. The pDockQ score can be linked to a positive predictive value (PPV), estimating the probability that the solution is a true positive (Das et al., 2026).

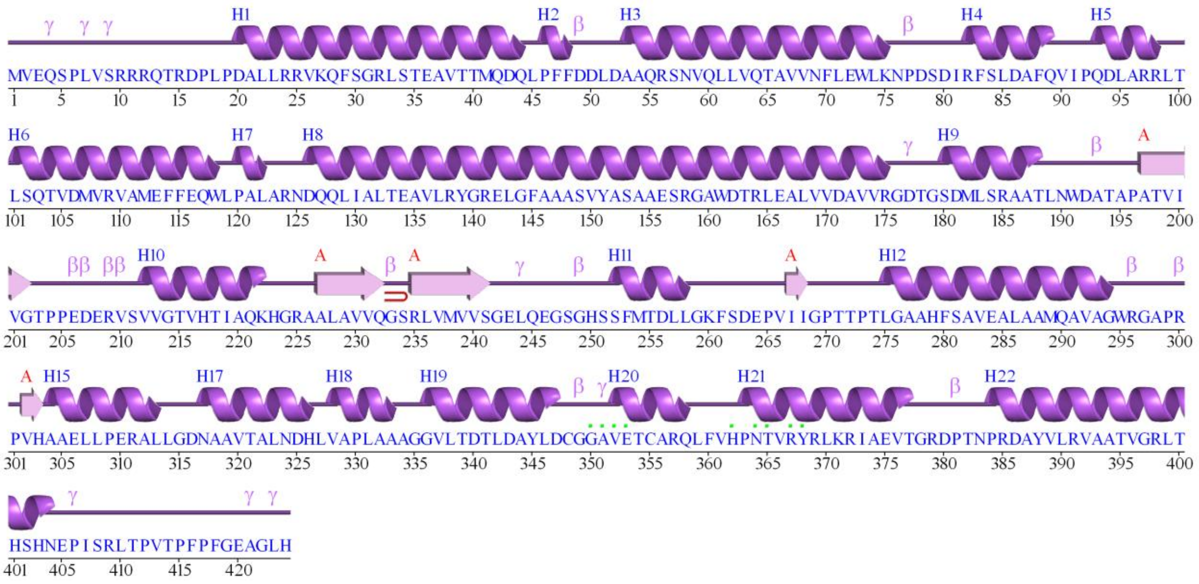


**Supplementary Figure S4**. PDBsum data of $\text{MabR}_{RO}$ using the structure prediction of dimer with DNA. Motifs: β *beta turn*; γ gamma turn; beta hairpin. Residue contacts: to DNA/RNA (Laskowski et al., 2018).

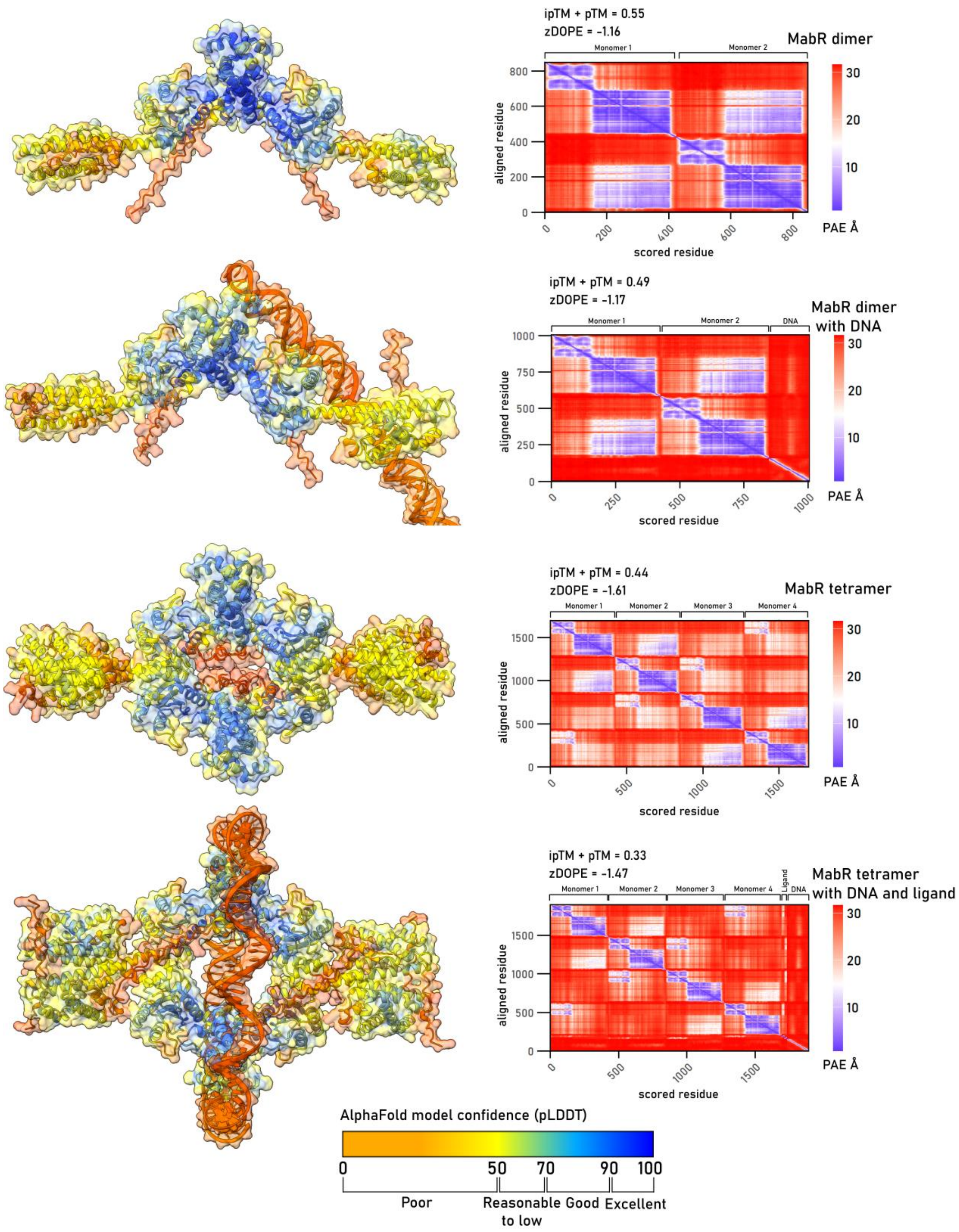


**Supplementary Figure S5**. Legend on next page.

**Supplementary Figure S5**. Cartoon representations of the models generated by AlphaFold3 (MabR) in dimeric and tetrameric formation, with or without ligand. The predictions are coloured according to the predicted local distance difference test (pLDDT), which indicates the local quality of the model, as shown in the legend. For each structure, the predicted aligned error (PAE), normalised discrete optimised protein energy (zDOPE), and overall predicted template modelling (pTM) scores are displayed. The PAE predicts the positional error for each residue x (scored residue) in a protein structure when aligned with its corresponding residue y (aligned residue) in the true structure. A zDOPE score below -1 suggests that the atom pair distances in the model are similar to those in a large sample of known protein structures, with at least 80% of the model's Cα atoms within 3.5 Å of their correct positions. As all the predicted structure are multimers, the pDockQ and AlphaFold-Multimer model confidence (0.8ipTM + 0.2pTM). The pTM score (ranging from 0 to 1) is the predicted TM score for a superposition between the predicted and the hypothetical true structure, indicating the accuracy of the prediction within a single chain. Contrary to the interface pTM (ipTM, ranging from 0 to 1), that evaluates the similarity between the true and predicted structures over interfacing residues, reflecting the accuracy of the prediction for a complex. The pDockQ score (ranging from 0 to 1) is another confidence criterion for protein complexes, considering the number of interfacing residues and their pLDDT score. The pDockQ score can be linked to a positive predictive value (PPV), estimating the probability that the solution is a true positive (Das et al., 2026).

A

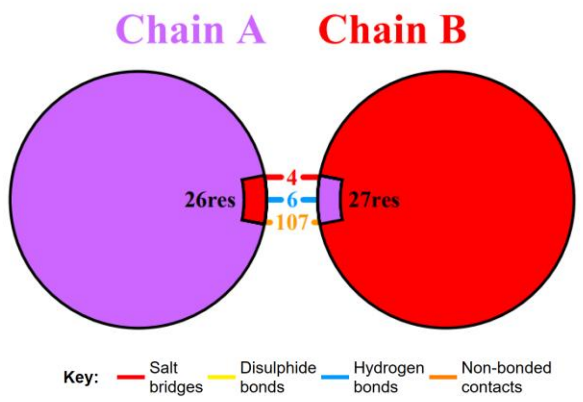


B

### Interface statistics

| Chain | No. of interface residues | Interface area (Å$^2$) | No. of salt bridges | No. of disulphide bonds | No. of hydrogen bonds | No. of non-bonded contacts |
|---|---|---|---|---|---|---|
| A | 26 | 1582 | 4 | - | 6 | 107 |
| B | 27 | 1570 | | | | |

C

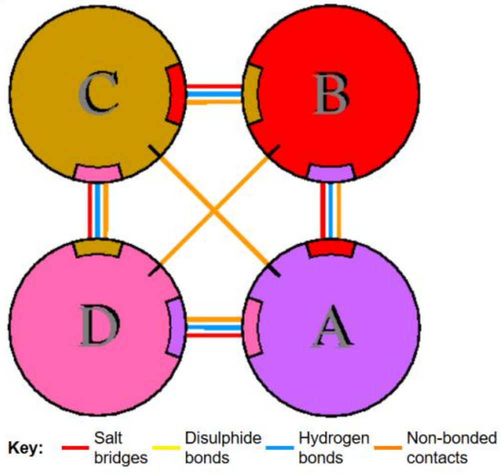


D

### Interface statistics

| Chains | No. of interface residues | Interface area (Å$^2$) | No. of salt bridges | No. of disulphide bonds | No. of hydrogen bonds | No. of non-bonded contacts |
|---|---|---|---|---|---|---|
| A-B | 38 : 38 | 2285 : 2278 | 4 | - | 8 | 194 |
| & C-D | 41 : 39 | 2270 : 2292 | 4 | - | 12 | 189 |
| A-C | 1 : 1 | 65 : 62 | - | - | - | 6 |
| & B-D | 1 : 1 | 37 : 38 | - | - | - | 4 |
| A-D | 45 : 47 | 2819 : 2830 | 6 | - | 9 | 258 |
| & B-C | 48 : 47 | 2870 : 2877 | 6 | - | 9 | 265 |

**Note**. Indented interfaces in the table are equivalent to the last prior non-indented interface. Equivalent chains are listed below.

**Equivalent chains:** A = B = C = D

**Supplementary Figure S6**. PDBsum analysis (Laskowski et al., 2018).

**Supplementary Table S1**. Overview of oligonucleotide sequences.

| Oligo-nucleotide name | Sequence (5’ - 3’) | Purpose |
|---|---|---|
| PL_0003 | TAAGAAGGAGATATACATATGACTATGGTCGAGCAGA | Pick up *mabR* with GA overhangs (FW) |
| PL_0004 | GTGGTGGTGGTGGTGGTGCAGCCCTGCTTCTCCGAATGGGAAGGG | Pick up *mabR* with GA overhangs (RV) |
| PL_0007 | TAAGAAGGAGATATACATATGGCGGTTCCGAAGGCGTCGG | Pick up *fadR1*$_{RO}$ with GA overhangs (FW) |
| PL_0008 | GTGGTGGTGGTGGTGGTGGTCGTCGAGCGGAATGTCGGTGCGCTC | Pick up *fadR1*$_{RO}$ with GA overhangs (RV) |
| PL_0048 | TTAACACCAGTCACACCCTTCCC | Pick up EMSA fragment (291 bp) (FW) |
| PL_0049 | TCGAGGCCGGCAGCCTTCGA | Pick up EMSA fragment (291 bp) (RV) |
| PL_0077 | GTCGCGTCAAACAGTTCTCC | RT-qPCR primer *mabR* (FW) |
| PL_0078 | ACTCGAGGAAGTTGACGACC | RT-qPCR primer *mabR* (RV) |
| PL_0079 | AGGGTTCATCGACGACAAGG | RT-qPCR primer *fadR1*$_{RO}$ (FW) |
| PL_0080 | GAAATCCTTGATCGACTCGGC | RT-qPCR primer *fadR1*$_{RO}$ (RV) |
| PL_0081 | GATCTGCTCGACCGTCTGA | RT-qPCR primer *atpClpX* (FW) |
| PL_0082 | GCGGATCATGATGCACCAG | RT-qPCR primer *atpClpX* (RV) |
| PL_0091 | GTGGTGGTGGTGGTGGTGTGTGTCCTTCTCCGAATCGTGTTG | Pick up *fadR2*$_{RO}$ with GA overhangs (FW) |
| PL_0092 | TAAGAAGGAGATATACATATGTTCGAGCCGATCGCACGGAAATCCGCCTCC | Pick up *fadR2*$_{RO}$ with GA overhangs (RV) |
| PL_0101 | TTCCACACCCGGTTCATGG | RT-qPCR primer *fabD* (FW) |
| PL_0102 | CCAGTTTAGTCAGTGCGTCG | RT-qPCR primer *fabD* (RV) |
| PL_0103 | CAAGTACGGCGTGAAGATCC | RT-qPCR primer *acpM* (FW) |
| PL_0104 | TCACTCCTCGTCGTTCTTGG | RT-qPCR primer *acpM* (RV) |
| PL_0105 | TTCCCCAACATCGTCGTCAC | RT-qPCR primer *kasAB* (FW) |
| PL_0106 | TCACCGGAAGATCGTTCTCC | RT-qPCR primer *kasAB* (RV) |
| PL_0107 | CACAGATTTCCGTCGTCCTC | RT-qPCR primer *accD6* (FW) |
| PL_0108 | ACCATGTCGACCTGCTCAC | RT-qPCR primer *accD6* (RV) |
| PL_0110 | GTGCAATTTTGTGGGAACCCAACAAATCCGCCGACAAAGGTTCATCCGCAACGACATCTCGCGGCACGCGAGGCTGGTGTCAAAAGGTGAGG | Pick up part 1 (Promoter + mCherry) from *pDD267* plasmid with GA overhangs (FW) |

| | | |
|---|---|---|
| PL_0111 | CGGTATCAGCTCACTCAAAGGCTCACTTATACAACTCATCCAT | Pick up part 2 (mCherry + backbone) from *pDD267* plasmid with GA overhangs (RV) |
| PL_0112 | TGGCATGGATGAGTTGTATAAGTGAGCCTTTGAGTGAGCTGATACC | Pick up part 2 (mCherry + backbone) from *pDD267* plasmid with GA overhangs (FW) |
| PL_0113 | ATTTGTTGGGTTCCCACAAAATTGCACGTGGGGTCCTCGCATCGAATCAGACCTGAATGGCGAATGGACGCGCC | Pick up part 3 (backbone + promoter) from *pDD267* plasmid with GA overhangs (RV) |
| PL_0114 | CATATGTCTGACCAGGGAAAATAGCC | Pick up mutated promoter fragment (FW) |
| PL_0115 | AAGCGCATAAACTCTTTAATAATAGCC | Pick up mutated promoter fragment (RV) |
| PL_0126 | ACCGACCTGTATTACCAGCTG | RT-qPCR primer $fadR2_{RO}$ (FW) |
| PL_0127 | GTCCTTCTCCGAATCGTGTTG | RT-qPCR primer $fadR2_{RO}$ (RV) |
| EMSA-probe | TTAACACCAGTCACACCCTTCCCATTCGGAGAAGCAGGGCTGTAACGAACGTCTGTCCAGATTCGATGCGAGGACCCCACGTGCAATTTTGTGGGAACCCAACAAATCCGCCGACAAAGGTTCATCCGCAACGACATCTCGCGGCACGCCGTTGACGGTGTTCTCTTGAAGAGTGATTTCGTTGCTCGCCCCGGGTCAGGGCTCTCAGACTCCCGGCATGCTCATCCCTTGGTTGGAGCTGCCGGGTTCTCATGACCGTGTCGCACTGTGGTCGAAGGCTGCCGGCCTCGA | 292 bp DNA probe used for EMSA |

**Supplementary Table S2**. Statistical analysis of $log_2$ fold change ($log_2FC$) values from RT-qPCR analyses. For each condition and gene, the 95% confidence interval (CI) of the mean $log_2FC$ and the p-value from a one-sample, two-tailed Student's t-test ($H_0$: $log_2FC = 0$) are reported. Conclusions indicate whether expression changes were statistically significant relative to baseline (CI including values between 0 and ±1) or reflected biologically relevant up- or downregulation (CI lower bound > 1 or CI upper bound < −1, respectively).

| **Conditions** | **Gene** | **95% Confidence interval (CI) of $Log_2FC$** | **Result T-test (p-value)** | **Conclusion** |
|---|---|---|---|---|
| $C_{16}$ fatty acid **3 hours** | *fabD* | [-3.08, 1.38] | 0.1298 | |
| | *acpM* | [-1.66, 1.70] | 0.9008 | |
| | *accD6* | [-3.48, 3.42] | 0.9221 | |
| $C_{18}$ fatty acid **3 hours** | *fabD* | [-3.11, -0.12] | 0.0433 | Differential expression |
| | *acpM* | [-1.08, 1.41] | 0.6283 | |
| | *accD6* | [-0.52, 0.17] | 0.1607 | |
| $C_{20}$ fatty acid **3 hours** | *fabD* | [-3.10, 1.22] | 0.2008 | |
| | *acpM* | [-4.62, 4.56] | 0.9814 | |
| | *accD6* | [-2.14, 2.10] | 0.9741 | |
| $C_{22}$ fatty acid **3 hours** | *fabD* | [-2.49, 0.57] | 0.1141 | |
| | *acpM* | [-1.24, -0.44] | 0.0120 | Differential expression |
| | *accD6* | [-1.09, 0.05] | 0.0596 | |
| $C_{16}$ fatty acid **16 hours** | *fabD* | [-5.73, 9.60] | 0.1920 | |
| | *acpM* | [-3.80, 6.17] | 0.2035 | |
| | *accD6* | [0.65, 1.31] | 0.0170 | Differential expression |
| $C_{18}$ fatty acid **16 hours** | *fabD* | [-1.92, 3.84] | 0.2879 | |
| | *acpM* | [-4.82, 4.65] | 0.9471 | |
| | *accD6* | [-1.62, 2.66] | 0.4044 | |
| $C_{20}$ fatty acid **16 hours** | *fabD* | [-0.87, 1.92] | 0.2484 | |
| | *acpM* | [-2.83, 2.49] | 0.8088 | |
| | *accD6* | [-1.87, 1.40] | 0.6020 | |
| $C_{22}$ fatty acid **16 hours** | *fabD* | [-1.43, 5.26] | 0.0665 | |
| | *acpM* | [-2.54, 1.61] | 0.4383 | |
| | *accD6* | [-1.00, 0.45] | 0.2472 | |
| C/N 9.5 vs. C/N 38 | *fabD* | [0.44, 3.48] | 0.0301 | Differential expression |
| | *acpM* | [-0.47, 4.92] | 0.070 | |
| | *kasAB* | [2.12, 2.62] | 0.0006 | **Upregulation** |
| | *accD6* | [0.53, 3.99] | 0.0302 | Differential expression |
| C/N 9.5 vs. C/N 184 | *fabD* | [2.55, 2.96] | 0.0003 | **Upregulation** |
| | *acpM* | [1.77, 5.15] | 0.0126 | **Upregulation** |
| | *kasAB* | [2.05, 3.51] | 0.0036 | **Upregulation** |

| | | | | |
|---|---|---|---|---|
| | *accD6* | [2.21, 4.03] | 0.0046 | **Upregulation** |
| pH 7 vs. pH 5.5 | *fabD* | [-1.09, 2.47] | 0.1278 | |
| | *acpM* | [-4.68, -0.27] | 0.0445 | Differential expression |
| | *kasAB* | [-7.29, 8.75] | 0.4529 | |
| | *accD6* | [-2.54, 5.96] | 0.1229 | |
| pH 7 vs. pH 8 | *fabD* | [-0.30, 1.55] | 0.0737 | |
| | *acpM* | [-10.97, 13.03] | 0.4732 | |
| | *kasAB* | [-7.25, 8.69] | 0.4582 | |
| | *accD6* | [-3.69, 2.84] | 0.3443 | |
| 29 °C vs. 22 °C | *fabD* | [-1.61, 1.61] | 0.9989 | |
| | *acpM* | [-9.92, 0.85] | 0.0685 | |
| | *kasAB* | [-2.65, 1.30] | 0.2801 | |
| | *accD6* | [-0.81, 1.19] | 0.4934 | |
| 29 °C vs. 34 °C | *fabD* | [-1.68, 3.96] | 0.2249 | |
| | *acpM* | [-8.55, -2.21] | 0.018 | **Downregulation** |
| | *kasAB* | [-0.95, 1.64] | 0.3737 | |
| | *accD6* | [0.76, 1.77] | 0.0084 | Differential expression |

**Supplementary Table S3.** Potential FadR-binding sites were identified in the upstream regions of *R. opacus* PD630 genes homologous to *M. tuberculosis* genes that exhibited altered expression following INH and ETH treatment. The search for FadR-binding sites in the upstream regions of these genes was based on the $FadR_{MT}$-binding consensus sequence described by Biswas et al., (2013). The binding sites, shown in the 5' to 3' direction, are present at a particular distance from the start codon. If available, the function of the gene is also provided.

| Location | NCBI identifier | Annotation | Distance to ORF | Predicted binding site | P-value |
|---|---|---|---|---|---|
| Pd630_LPD06939 | AHK34124 | FadR regulatory protein paralog 1 | 81 bp | ATTA**GGT**CTGA**CC**GTTA | |
| Pd630_LPD05300 | AHK32505 | *fabD* (Fatty acid synthase II operon) | 81 bp | C**A**AA**GGT**TCAT**CC**GCAA | |
| Pd630_LPD05301 | AHK32506 | *acpM* meromycolate extension acyl carrier protein | -107 bp | T**A**CC**GGT**GACCT**C**CTCG | |
| Pd630_LPD05140 | AHK32346 | FadR regulatory protein paralog 2 | 35 bp | C**A**CT**GGT**AAGA**CC**ACTT | |
| Pd630_LPD07419 | AHK34604 | FadR regulatory protein paralog 3 | 82 bp | A**A**CA**GGT**ACAC**CC**GCAC (reverse complementary) | |
| Pd630_LPD00455 | AHK27699 | Diacylglycerol O-acyltransferase | -67 bp | C**A**CT**GGT**ACTG**CC**TGTG | 0.000152 |
| Pd630_LPD00515 | AHK27759 | Putative cysteine desulferase | 75 bp | CGAA**GGT**GCGT**CC**GATT | 0.005803 |
| Pd630_LPD06301 (1) | AHK33488 | Isocitrate lyase | -206 bp | T**A**CG**GGT**GTGC**C**GCGGC (reverse complementary) | |
| Pd630_LPD06301 (2) | AHK33488 | Isocitrate lyase | 334 bp | G**A**TT**GGT**GATTG**C**AGTC | |

| | | | | | |
|---|---|---|---|---|---|
| Pd630_LPD06294 | AHK33481 | Hypothetical protein (anti-sigma B factor antagonist) | -6 bp | A**A**CA**GGT**CGAT**C**TCGCC (reverse complementary) | |
| Pd630_LPD05154 | AHK32361 | Hypothetical protein | 17 bp | G**A**CC**GGT**TGCA**C**GTTCA | |
| Pd630_LPD04240 | AHK31453 | 2-haloacid dehalogenase | -89 bp | C**A**CC**GGT**ATCG**CC**GCCG | |
| Pd630_LPD01228 (1) | AHK28461 | putative inactive (triacylglycerol) lipase | 66 bp | G**A**CG**GGT**ATCG**C**AATAC | 0.002190 |
| Pd630_LPD01228 (2) | AHK28461 | putative inactive (triacylglycerol) lipase | 277 bp | CGTGC**G**GGTGA**CC**GCTT | 0.002190 |
| Pd630_LPD05220 (1) | AHK32720 | Hypothetical protein | 98 bp | A**A**GA**GGT**GGTGA**C**CCGC (reverse complementary) | |
| Pd630_LPD05220 (2) | AHK32720 | Hypothetical protein | 178 bp | C**A**TG**GGT**GGACA**C**AAAC | |
| Pd630_LPD01727 | AHK28956 | Uncharacterised protein | -111 bp | G**A**GA**GGT**ACGCG**C**CGCG | |
| Pd630_LPD05549 | AHK32747 | Putative sterigmatocystin biosynthesis fatty acid synthase subunit alpha (FASI) | 147 bp | C**A**CC**GGT**CACTT**C**GCGG | |
| Pd630_LPD04067 | AHK31280 | Universal stress protein | -279 bp | C**A**CG**GGT**CGGC**C**GTGCC | |
| Pd630_LPD03066 | AHK30288 | acyl-CoA dehydrogenase | 311 bp | G**A**TC**GGT**TCGC**C**GAACG (reverse complementary) | |

| | | | | |
|---|---|---|---|---|
| Pd630_LPD00768 | AHK28008 | Hypothetical protein | 74 bp | A**A**CG**GGT**GCGT**CC**A CGC |

**Supplementary Table S4.**Genome wide predicted binding sites MabR and corresponding downstream annotated genes. Genes were annotated by identifying open reading frames (ORFs) in proximity to the binding sites and performing BLAST searches against *Rhodococcus* sequences (Altschul et al., 1990).

| Potential binding site MabR | Function of gene(s) | Accession number | Number |
|---|---|---|---|
| TTTTGT(N)$_9$ACAAAT | Fatty acid synthase complex II (FASII) | Pd630_LPD05300 | 1 |
| TTTTGT(N)$_9$ACAA**CG** | Putative ABC transporter ATP-binding protein YdiF | Pd630_LPD01538 | 2 |
| TTTTGT(N)$_9$**C**CA**G**AA | Acetolactate synthase | Pd630_LPD03152 | 3 |
| TTTTGT(N)$_9$**G**CAAA**C** | TerD family protein; stress response | Pd630_LPD06671 (part 1) Pd630_LPD06670 (part 2) | 4 |
| TTTTGT(N)$_9$A**G**AAA**C** | Sugar porter family MFS transporter | WP_005250354.1 Pd630_LPD07294 Pd630_LPD07295 | 5 |
| TTTTGT(N)$_9$ACA**GG**A | Carbohydrate diacid regulator (CdaR) | Pd630_LPD07475 | 6 |
| TTT**C**GT(N)$_9$ACAAAA | Carbon monoxide dehydrogenase operon | Pd630_LPD07873 Pd630_LPD07874 Pd630_LPD07875 Pd630_LPD07876 | 7 |
| T**G**TTGT(N)$_9$ACAAAT | Alkene reductase | Pd630_LPD07077 | 8 |
